# Behavioral estimates of mating success corroborate genetic evidence for pre-copulatory sexual selection in male *Anolis sagrei* lizards

**DOI:** 10.1101/2023.02.04.527135

**Authors:** Rachana S. Bhave, Heidi A. Seears, Aaron M. Reedy, Tyler N. Wittman, Christopher D. Robinson, Robert M. Cox

## Abstract

In promiscuous species, fitness estimates obtained from genetic parentage may often reflect both pre- and post-copulatory components of sexual selection. Directly observing copulations can help isolate the role of pre-copulatory selection, but such behavioral data are difficult to obtain in the wild and may also overlook post-copulatory factors that alter the relationship between mating success and reproductive success. To overcome these limitations, we combined genetic parentage analysis with behavioral estimates of size-specific mating in a wild population of brown anole lizards (*Anolis sagrei*). Males of this species are twice as large as females and multiple mating among females is common, suggesting the scope for both pre- and post-copulatory processes to shape sexual selection on male body size. Our genetic estimates of reproductive success revealed strong positive directional selection for male size, which was also strongly associated with the number of mates inferred from parentage. In contrast, a male’s size was not associated with the fecundity of his mates or his competitive fertilization success. By simultaneously tracking copulations in the wild via the transfer of colored powder to females by males from different size quartiles, we independently confirmed that large males were more likely than small males to mate. We conclude that body size is primarily under pre-copulatory sexual selection in brown anoles, and that post-copulatory processes do not substantially alter this pre-copulatory selection. Our study also illustrates the utility of combining both behavioral and genetic methods to estimate mating success to disentangle pre- and post-copulatory processes in promiscuous species.

## Introduction

In species where females mate promiscuously with multiple partners, sexual selection on male traits can continue to occur after copulation, through sperm competition and female cryptic choice. These post-copulatory processes can alter the siring success of males and thereby modify the strength of sexual selection on traits linked to mating success (Parker 1970; Kvarnemo and Simmons 2013; Simmons et al. 2017; Glaudas et al. 2020). For example, larger males may mate with more females, but this may not translate into strong sexual selection if they are poor sperm competitors. In addition to pre- and post-copulatory sexual selection, the net reproductive fitness of a male is also influenced by the fecundity of his mating partners (Wong and Candolin 2005; Venner et al. 2010; Pincheira-Donoso and Hunt 2017). For any given trait, total selection due to variance in reproductive success can thus be partitioned into selection acting through variance in pre-copulatory mating success, post-copulatory fertilization success, and female fecundity (Arnold and Wade 1984; Koenig et al. 1991; Collet et al. 2012; Pélissié et al. 2014).

Furthermore, selection mediated through any one of these components of fitness may be reinforced or weakened by selection acting through other components (Arnold and Wade 1984; Shuster et al. 2013). Therefore, a complete understanding of selection on a given trait requires estimating phenotypic selection as a function of total reproductive success as well as its underlying components (Arnold and Wade 1984). However, our ability to partition sexual selection in wild populations is hindered by both the cryptic nature of post-copulatory processes and the difficulty of independently measuring mating success, mate fecundity, and fertilization success (Droge-Young et al. 2012; Oneal and Knowles 2015; Marie-Orleach et al. 2016).

Studies of sexual selection in wild populations have typically measured fitness using either genetic estimates of parentage or behavioral observations of mating success. However, either of these approaches can provide an incomplete picture of sexual selection when considered alone (Thompson et al. 2011; Marie-Orleach et al. 2016; Olsson et al. 2019), making it difficult to disentangle pre- and post-copulatory selection (Danielsson 2001; Mobley and Jones 2013; Kamath and Losos 2018; Cramer et al. 2020). Genetic parentage analysis can identify mating pairs from shared parentage, thereby providing minimum estimates of the number of mating partners for both males and females (Flanagan and Jones 2019). Such data can be used to estimate both pre-copulatory mating success (minimum number of known mates per male) and post-copulatory fertilization success (proportion of offspring sired with females who have multiple known mates) (Rose et al. 2013; Evans and Garcia-Gonzalez 2016). However, many copulations may go undetected if females mate with many partners but produce relatively few offspring, potentially leading to mis-estimation of selection via mating success. (Pemberton et al. 1992; Flanagan and Jones 2019; Olsson et al. 2019; Baird and York 2021). Direct observations of copulations avoid this problem, but it is usually impossible to comprehensively track all copulations in wild populations. For example, animals may copulate in obscure or sheltered locations, the duration of mating may be short, the population density may be too low, or the population size may be too high for comprehensive observations (Candolin 1998; Dunn et al. 2012; Johnson et al. 2014; Cramer et al. 2020). Therefore, reliance on either genetic or behavioral methods alone to measure fitness may lead to misestimation of the strength of pre- and post-copulatory sexual selection (Pischedda and Rice 2012; Evans and Garcia-Gonzalez 2016; Baird and York 2021). Consequently, there is increasing emphasis on approaches that measure fitness and its components using a combination of both behavioral observations and genetic parentage analyses to help partition pre- and post-copulatory dimensions of sexual selection (Collet et al. 2012; Pischedda and Rice 2012; Evans and Garcia-Gonzalez 2016; McDonald et al. 2017; Simmons et al. 2017; Olsson et al. 2019).

We studied the sexually dimorphic brown anole lizard, *Anolis sagrei*, to determine which components of male reproductive success (i.e., mating success, average mate fecundity, or competitive fertilization success) generate selection for larger body size in this species. Adult male brown anoles are, on average, two to three times larger than adult females in body mass (Cox and Calsbeek 2010a). Larger males are more likely to succeed in competitive interactions that lead to female encounters and to sire more offspring (Tokarz 1985; Kamath and Losos 2018). However, female brown anoles produce offspring with multiple sires during the breeding season and can store sperm for several months (Calsbeek et al. 2007; Calsbeek and Bonneaud 2008; Duryea et al. 2016; Kamath and Losos 2018; Kahrl et al. 2021). Females may also bias their offspring sex ratio based on the body size or condition of the males with which they mate, suggesting that post-copulatory processes can also shape selection on male body size (Calsbeek and Bonneaud 2008; Cox and Calsbeek 2010b; Cox et al. 2011). Although several studies have detected selection for larger body size in male brown anoles (Cox and Calsbeek 2010a; Duryea et al. 2016; Kamath and Losos 2018), no study to date has assessed the extent to which the higher reproductive success of larger males is due to higher mating success, higher average mate fecundity, higher fertilization success, or a combination of these components of reproductive success (Friesen et al. 2020).

Given the scope for both pre- and post-copulatory selection to act on male body size in brown anoles (Calsbeek et al. 2007; Kahrl et al. 2016), we combined genetic parentage and behavioral observations of mating to estimate fitness components in a wild population of this species. Based on the established role of body size in mediating aggressive interactions among males, we hypothesized that body size is primarily subject to pre-copulatory selection (Tokarz 1985; Duryea et al. 2016; Kamath and Losos 2018) Specifically, we predicted that body size would be positively associated with both total reproductive success (number of offspring sired) and mating success (number of mates identified via genetic parentage). Although anoles only lay one egg at a time, larger females produce more offspring compared to smaller females and tend to be more fecund (Andrews & Rand, 1974; Cox & Calsbeek, 2011; Duryea et al., 2016; Warner & Lovern, 2014). Thus, we also explored whether large males preferentially mate with larger and more fecund females. Since post-copulatory selection could weaken or reinforce pre-copulatory selection (Danielsson 2001; Hosken et al. 2008; Kvarnemo and Simmons 2013; Parker et al. 2013; Turnell and Shaw 2015), we also tested whether competitive fertilization success (i.e., the proportion of offspring sired with females who also produced offspring with other males) differed as a function of male body size. To corroborate our inferences based on genetic parentage with behavioral estimates of mating success, we quantified size-specific mating rates in the wild by tracking the copulatory transfer of fluorescent powders from males to females, with different colors of powder corresponding to different quartiles for male body size. We then tested whether larger males obtained a greater number of copulations, whether larger males mated with larger females, and whether female body size and fecundity were positively correlated. Our study design thus allowed us to separate the contributions of pre-copulatory mating success, female fecundity, and post-copulatory fertilization success.

## Methods

### Field site and sampling

We studied an island population of brown anole lizards (*Anolis sagrei*) in the Guano Tolomato Matanzas Natural Estuarine Research Reserve in northern Florida (29°37′ ′′ 81°12′ 46′′ W). Adults begin mating around March (Lee et al. 1989) and females typically lay one egg every 7-14 days from April through October. Juveniles emerge between late May and November, and most do not enter the breeding population as adults until the subsequent year. To assay the reproductive success of males in the wild, we sampled all adults and juveniles of the population at four different times during the breeding season (March, May, July, and October) in 2019. We marked each new individual with a unique toe clip and preserved a small (1-2 cm) tail clip in 100% ethanol at -20°C for genotyping. We measured snout-vent length (SVL, nearest 1 mm) and body mass (nearest 0.01g) of all individuals prior to releasing them at their site of capture the following day. We captured and measured a total of 920 adults (hatched prior to 2019) and 905 juveniles (hatched in 2019) on the island. Most of the adults were first captured as hatchlings in their year of birth and genotyped during previous sampling censuses.

### Genotyping and parentage assignment

We extracted DNA by adding 3-5 mg of tail tissue to 150 µl of 10% Chelex^R^ resin (Bio-Rad, Inc.) with 1.4 µl of Proteinase K (20 mg/ml, Qiagen, Chatsworth, CA), incubating at 55°C for 180 min, and denaturing at 99°C for 10 min. If the DNA concentration was not within the desired range of 10-25 ng/µl, we repeated extractions and modified the above protocol by incubating new tail samples in 40 µl of 10% Chelex^R^ resin with 1.5 µl of Proteinase K. After centrifugation at 2250 *g* at 4°C for 15 minutes, we collected 3 µl of supernatant from these extractions to genotype individuals using the Genotyping-in-Thousands by sequencing (GT-seq) protocol (Campbell et al. 2015) with a custom panel of primers for 215 biallelic SNP loci that were previously identified from RAD-seq data (HA Seears, unpublished). For all extractions with an average DNA concentration of <10 ng/µl (*n* = 282 of 1319 samples), we carried out an additional purification step on the supernatant using 1.8x volume of AMPure XP beads (Beckman Coulter, Brea, CA, USA) and eluted samples in 20 µl 1x TE (Fisher Bioreagents, Fair Lawns, NJ, USA) to concentrate the DNA to >10 ng/µl. After extraction, we shipped DNA samples to GTseek LLC (Twin Falls, ID, USA) for library preparation, sequencing, and data processing to obtain genotypes. Briefly, all 215 loci were simultaneously amplified and tagged with Illumina priming sequences in a multiplexed polymerase chain reaction (PCR). Each sample was then tagged with well-specific and plate-specific indices in a second PCR. The PCR products were then standardized to similar concentrations, pooled, cleaned, and then sequenced on an Illumina NextSeq 550 with 1 × 75 bp reads. The raw Illumina reads were checked for quality using FastQC and then de-multiplexed and assigned genotypes following Campbell et al. (2015).

We used SNPPIT 2.0 (Anderson 2012) to assign genetic parentage. We included all offspring known to have hatched in 2019 that were successfully genotyped at a minimum of 128 loci (< 40% missing loci; *n* = 885 successfully genotyped of 905 total offspring). We included adults as potential parents if they were successfully genotyped at a minimum of 165 loci (< 23% missing loci). Since individuals that were present but were not captured in 2019 may have also produced offspring in that year, we included all successfully genotyped individuals captured on the island between 2015 and 2018 as potential parents (*n* = 7042 individuals genotyped in previous studies). Of these putative parents, 870 individuals were captured as adults in 2019. We used a significance threshold of *P* < 0.05 after correcting for the false discovery rate (FDR) to assign parentage. We successfully assigned 736 offspring (83.2% of 885 genotyped offspring) to a total of 610 parents (*n* = 357 females, 253 males). Of these 610 parents, 479 (78.5%) were among the 870 successfully genotyped adults that we captured in 2019 (*n* = 276 females, 203 males) and 131 (21.4% of 610) were only captured in previous sampling years (*n* = 81 females, 50 males). Because we did not measure body size for this subset of 50 adult males in 2019, we excluded them from our calculations of relative fitness and our analyses of sexual selection.

Among the 870 successfully genotyped adults that we captured in 2019, a total of 391 individuals (*n* = 213 females, 178 males, 44.9%) were found to have zero reproductive success, since they were included in the SNPPIT analysis but were not assigned offspring.

### Partitioning reproductive success and measuring selection

All statistical analyses were performed in R v. 4.2.1 (R Core Team 2022) using the RStudio interface (RStudio Team 2022). We conducted univariate selection analyses to test whether body size of males was a predictor of reproductive success (total number of offspring sired in 2019) and its components (i.e., mating success, average mate fecundity, and competitive fertilization success), as estimated by genetic parentage. We measured mating success as the total number of unique females with which a male sired offspring. We measured average mate fecundity as the mean number of offspring produced across all female partners of a male, including offspring sired by other males. We measured competitive fertilization success by calculating the mean proportion of offspring sired by a male with each of his partners. To detect competing males from parentage data, a female must produce at least two offspring that are assigned to at least two mates. Thus, our measure of competitive fertilization success excluded all situations in which females produced either a single offspring or multiple offspring sired by a single male (following Devigili et al. 2015). To account for the fact that the null expectation for proportional fertilization success decreases with the number of additional males with which a female has mated, we used the following formula (Devigili et al. 2015):

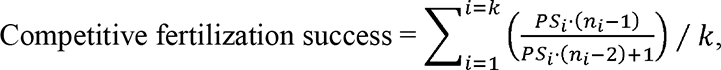

where be is the proportion of offspring sired for each female with which that male mated, w is the total number of females with which that male mated that had more than two mates, and, is the total number of mates of the female. Thus, a male that sired 33.3% of the offspring from a female that had three total mates would have a competitive fertilization success score of 0.5, which would be the same as a male that sired 50% of the offspring from a female that had only two mates.

We estimated univariate linear (*s*) and non-linear (*c*) selection differentials following Lande and Arnold (1983). We standardized body mass to a mean of 0 and a standard deviation of 1. We calculated relative fitness by dividing total reproductive success and each of its components (see above) by the mean value of that fitness component across all males in the population that were included in the analysis. We used ordinary least-squares regressions of each measure of relative fitness on standardized body mass to estimate univariate linear selection differentials, with separate regressions for each fitness component. To estimate *s*, we included only the linear term for body mass, and to estimate *c*, we included both the linear and quadratic terms (i.e., 0.5 × body mass^2^) (Lande and Arnold 1983; Stinchcombe et al. 2008). We used generalized linear models to test the significance of selection estimates. We used a negative binomial distribution for all components of fitness except competitive fertilization success, which had a Gaussian distribution. Non-linear selection differentials were not significant for any fitness component, so we only present visualizations of linear selection differentials. We considered individuals with zero reproductive success to have zero mating success, whereas the remaining fitness components were considered inestimable. This approach assumes that failure to reproduce is due to failure to mate when it could, in principle, also reflect low mate fecundity and/or poor competitive fertilization success. To confirm that including these zero values did not bias our partitioning of selection among components of reproductive success, we repeated the above analyses using only the subset of males that had at least one offspring (Fig. S1).

### Assessing size-specific mating success with fluorescent powders

We directly assessed the relationship between body size and copulation rates at two points in the middle of the breeding season: May 12-16 and July 26-Aug 3, 2019. In the first two days of each sampling period, we captured as many adult males on the island as possible and distributed them into size quartiles based on their body mass (May: *n* = 153; July: *n* = 128).

Before releasing each male to its site of capture the following day, we powdered males on their venters with one of four colors of fluorescent powder corresponding to their size quartiles (A/AX Series, DayGlo Color Corp., Ohio). The four colors (orange, yellow, pink and green) were selected after pilot studies confirmed that different colors of powder transferred during successive copulations could be clearly distinguished in the event of multiple mating across different size quartiles. These powders are non-toxic, easily differentiated under ultraviolet (UV) light, and wear off after a few days without negatively affecting the fitness of animals (Holbrook et al. 1970; Rojas-Araya et al. 2020). We switched the colors assigned to each size quartile between May and July to ensure that any observed mating patterns were not due to underlying differences in our ability to detect each color. We were not blind to the size quartile associated with the colors during the study. Subsequent studies in the same population using a double-blind study design have not uncovered significant biases in estimation of copulation rates (RS Bhave, unpublished). Two days after males were released, we captured as many adult females on the island as possible in a single day in May (*n* = 132) and across 5 days in July (*n* = 312; 50% of these captures occurred on the first day). We noted the color of any powder on or near the cloaca under UV light. Presence of color found on any other part of the body was uncommon and treated as a non-copulation contact. We tested whether observed copulations within each size quartile (as determined by the color of transferred powder) significantly differed from null expectation using a chi-square test with 3 degrees of freedom. The expected number of copulations for each size quartile was a product of the proportion of powdered males that were assigned to each quartile and the total number of copulations detected across all females.

We also used data on transfer of fluorescent powder to test whether large males mated more frequently with large females, potentially benefiting males through the increased fecundity of larger females. To test this prediction, we conducted an ordinary least square regression of female body mass (continuous dependent variable) on male size quartile (ordinal independent variable) and estimated significance with a type II ANOVA using the *car* package (Fox et al. 2019) in R. We tested the underlying assumption that female fecundity is positively correlated with body size by regressing the total number of offspring assigned to a female using genetic parentage (continuous response variable) on female body mass (continuous independent variable) using generalized linear models with a negative binomial error distribution and a logit link function. Because female body mass can vary depending on the presence or absence of oviductal eggs, we also repeated the above analyses by considering SVL as an alternate measure of female body size. In all cases, we conducted two separate analyses using data from May and July, followed by a third analysis on data combined across May and July.

### Comparing behavioral and genetic approaches

To compare behavioral and genetic approaches, we assessed whether males belonging to larger size quartiles in our powdering experiment (behavioral) also differed in their fitness components as measured by parentage (genetic). We conducted separate generalized linear regressions for males captured in May versus July, with reproductive success (negative binomial), mating success (negative binomial), average mate fecundity (negative binomial) and competitive fertilization success (Gaussian) as response variables. We treated the size quartile that males belonged to in each month as an ordinal predictor variable. In each analysis, we only considered males that were powdered in that month and successfully genotyped. To test whether associations between size quartile and fitness components varied across months, we repeated the above analyses on data combined across May and July while also including an effect of month and its interaction with size quartile. A subset of successfully genotyped males that were captured and powdered in May were also captured and powdered in July (*n* = 37), so these individuals were included twice in our combined analysis. Given that these males constituted only 15% of all individuals that were powdered, and that model results were similar with or without inclusion of these repeated measures across both months, we did not include individual ID as a random effect to simplify the model fit. We obtained effect sizes of all main effects in these models from a type II ANOVA unless the interaction of size quartile × month was significant, in which case we conducted a type III ANOVA.

We carried out a chi-square test with 3 degrees of freedom to test whether the number of copulations in each size quartile, as determined by powdering (observed), corresponded to the number of copulations predicted from genetic parentage (expected). To calculate the expected proportion of copulations in each size quartile, we first estimated the number of unique dam-sire pairs from genetic parentage for sires. We assumed that each parental pair indicates at least one copulation with a male belonging to a particular size quartile, then divided the total copulations assigned in each size quartile by the total number of copulations attributable to all males that were measured and powdered in either May or in July. The expected number of copulations was calculated by multiplying this proportion by the total number of copulations observed from the transfer of fluorescent powder in the respective months.

## Results

### Partitioning pre- and post-copulatory selection on body size

We found significant directional selection favoring large male body mass when using total reproductive success as a measure of fitness (*s* = 0.40 ± 0.08, χ = 22.43, *P* < 0.001, Fig. 1A), and we found similarly strong selection when using only its pre-copulatory component of mating success (*s* = 0.33 ± 0.07, χ = 19.36, *P* < 0.001, Fig. 1B). Directional selection favoring large size persisted when we excluded males who did not sire any progeny from our analyses using reproductive success and mating success (Fig. S1). However, neither average mate fecundity (*s* = -0.03 ± 0.04, χ = 0.15, *P* = 0.70, Fig. 1C) nor competitive fertilization success (adjusted for number of competing males) generated significant selection on male body mass (*s* = 0.02 ± 0.02, *F_1,115_* = 2.18, *P* = 0.17, Fig. 1D). There was no significant quadratic (non-linear) selection on male body mass with respect to total reproductive success (*c* = 0.30 ± 0.12, χ*^2^* = 2.51, *P =* 0.11), mating success (*c* = 0.30 ± 0.12, χ*^2^* = 2.33, *P =* 0.13), average mate fecundity (*c* = 0.19 ± 0.09, χ*^2^* =1.21, *P =* 0.27) or competitive fertilization success (*c* = - 0.0009 ± 0.02, *F_1,115_* = 0.001, *P =* 0.97).

**Fig. 1:**
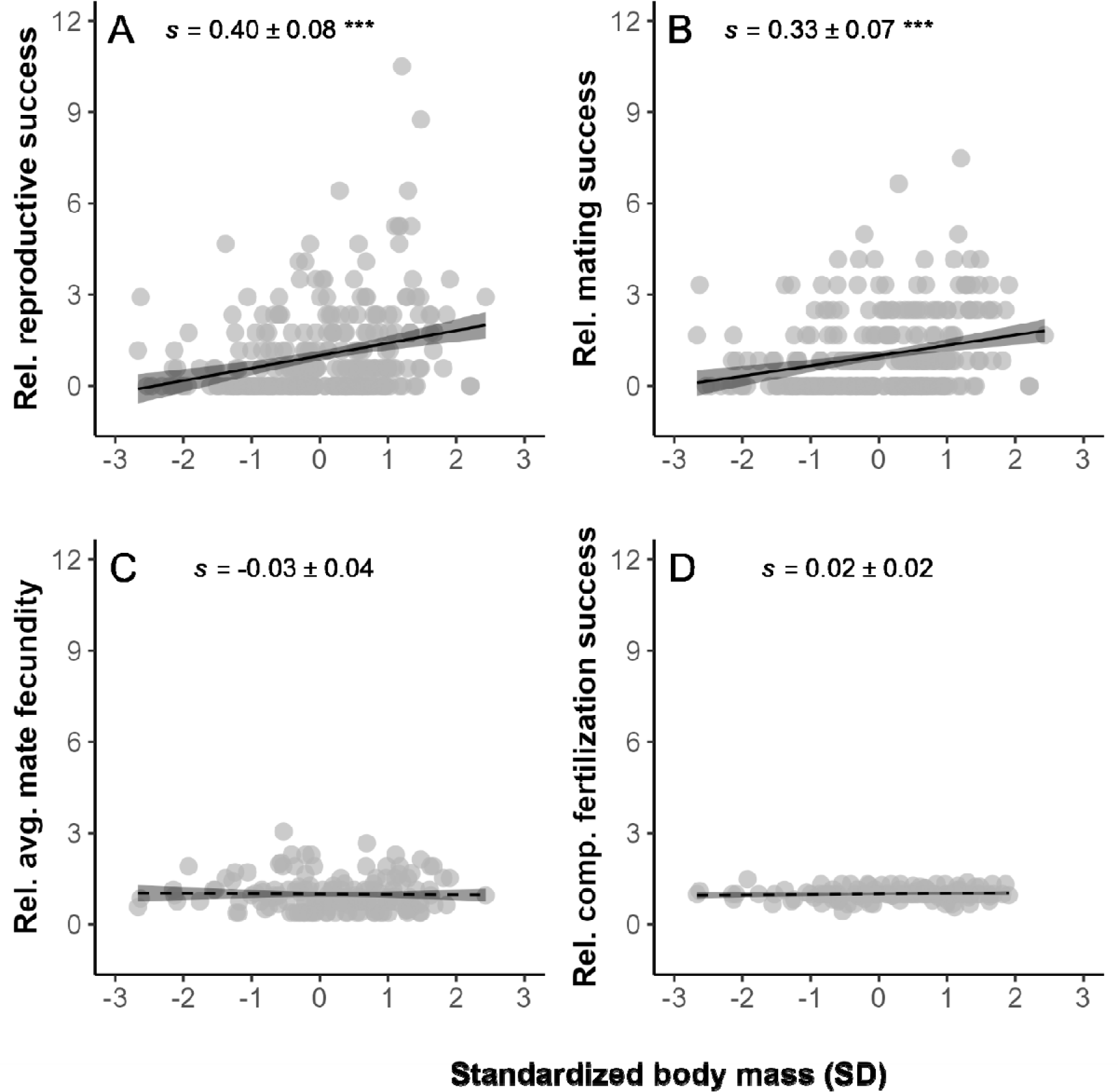
Linear selection on adult body mass as a function of different components of male fitness, including relative measures of (A) reproductive success (total number of offspring), (B) mating success (total number of mates), (C) average mate fecundity (mean fecundity across all mates), and (D) competitive fertilization success (mean proportion of offspring sired across all mates, adjusted for the number of competing males). For each fitness component estimated from genetic parentage, we divided individual fitness by the population mean to obtain relative measures. Adult body mass was measured at the start of the breeding season in March and standardized to a mean of 0 and standard deviation of 1. Trendlines with 95% confidence intervals (CI) from linear regressions are used to visualize linear selection. Solid lines and asterisks in panels A-B indicate significant selection differentials while dotted lines in panels C-D indicate non-significant selection differentials (*** *P* < 0.001).

### Behavioral estimates of size-specific mating success

We powdered a total of 241 males across May and July to test whether actual copulation rates differed across male size quartiles (Fig. 2A-C). Based on detection of transferred powder (Fig. 2D), we found that 38 of 132 (28.8%) females in May and 151 of 312 (48.4%) females in July mated within the five-day collection period, with most of these copulations occurring within three days of the release of powdered males. We also found that 1 of 38 (2.6%) females in May and 10 of 151 (6.6%) females in July mated with males from more than one size quartile during that period. We omitted 2 of 39 and 5 of 161 total copulations in May and July respectively, since we could not accurately resolve the color of fluorescent powder. Omitting these instances from the analyses did not bias the number of copulations for any size quartile. Within each month, observed copulations differed significantly from our null expectation of an equal number of matings across size quartiles (May: χ = 8.03, df = 3, *P =* 0.045, *n* = 37 copulations, July: χ = 8.33, df = 3, *P =* 0.039, *n* = 156 copulations; Fig. 2E-F). This difference was primarily attributable to the smallest size quartile having consistently fewer copulations than expected in each month. We saw a similar relationship between male size quartile and mating success after pooling data from both months (χ = 11.64, df = 3, *P =* 0.009, *n* = 193 copulations).

**Fig. 2:**
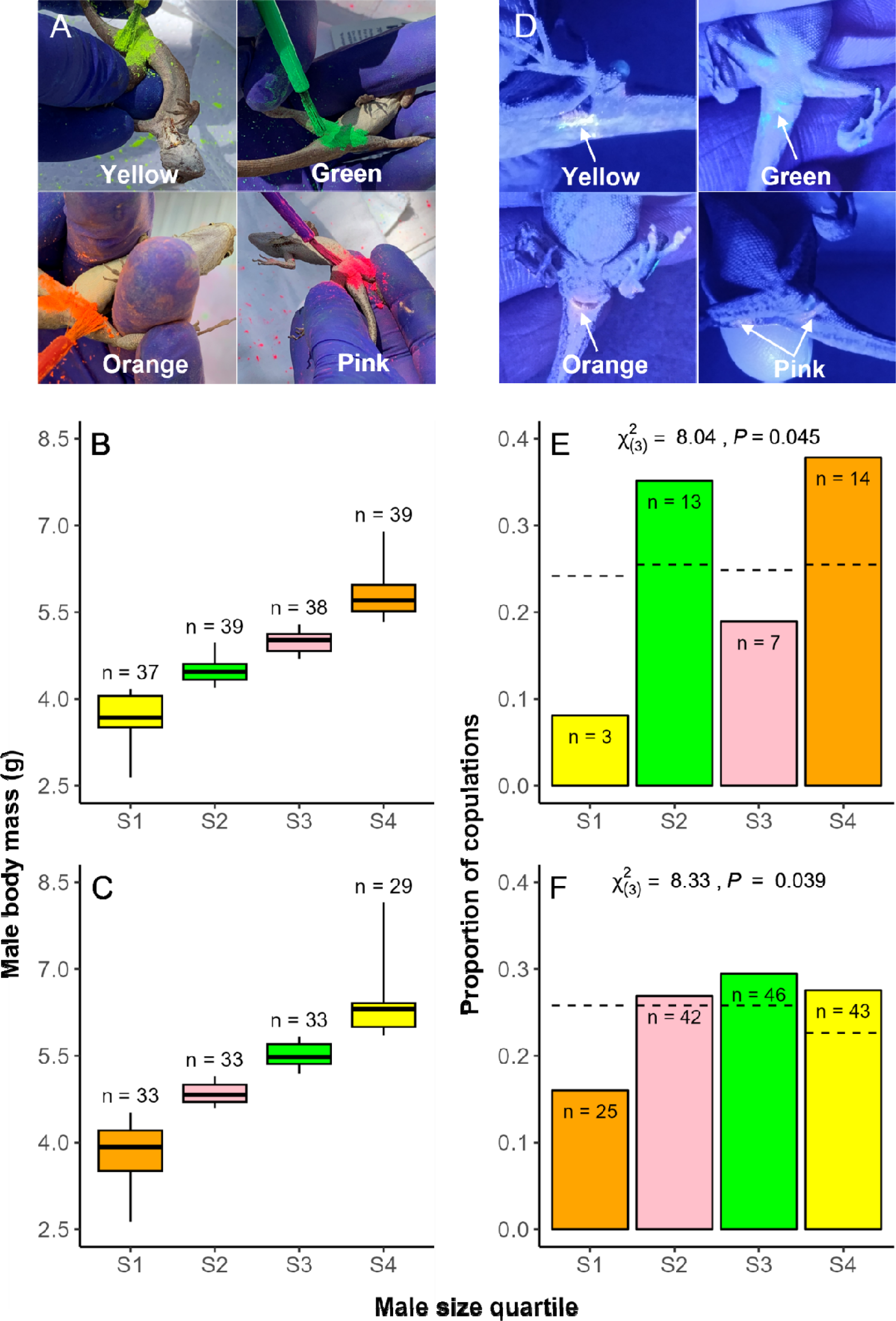
Procedure for detecting copulations in the wild. (A) Males were dusted with one of four colors of fluorescent powder based on size quartiles for body mass in (B) May, and (C) July, with colors alternated among size classes between months. Boxplots in B and C depict medians (lines), interquartile ranges (boxes), and minimum and maximum values (whiskers), with the number of males in each quartile shown above each boxplot. After males were released and allowed to interact freely with females for two days, females were captured and (D) inspected under UV light for the presence and color of any powder transferred near their cloaca. The proportions of total copulations detected among females that we correctly attributed to males from each size category are shown separately for (E) May, and (F) July. The number of females with each color of powder is indicated within each bar. The dotted lines give the expected proportion of copulations in each size quartile if mating is random with respect to male size. Colors of bars and box plots indicate the color of powder used for that size quartile.

Although there was a weak trend towards positive size-assortative mating, female body mass did not differ significantly across male size quartiles in either May (*F_3,33_* = 2.40, *P =* 0.085, Fig. 3A) or July (*F_1,152_* = 2.12, *P =* 0.10, Fig. 3B). However, when considering data combined across both months, we found a weak but significant positive correlation between female body mass and the size quartiles of males with which they mated (Size Quartile: *F_3,185_* = 3.24, *P* = 0.023, Month: *F_1,185_* = 3.71, *P* = 0.056, Size Quartile x Month: *F_3,185_* = 0.75, *P* = 0.52). Female SVL did not differ significantly across male size quartiles in May (*F_3,33_* = 1.48, *P =* 0.22, Fig. 3C), July (*F_1,152_* = 2.26, *P =* 0.083, Fig. 3D), or when combining both months (Size Quartile: *F_3,185_* = 1.48, *P* = 0.22, Month: *F_1,185_* = 1.23, *P* = 0.27, Size Quartile x Month: *F_3,185_* = 1.43, *P* = 0.24). The total number of offspring produced by a female in a year tended to increase with her body mass, but this weak relationship was not significant in May (X^2^ = 3.50, *P =* 0.061) or July χ χ = 2.54, *P =* 0.11). However, when considering SVL as a measure of female size, total number of offspring had a strong positive association with female body size in May (X^2^ = 9.91, *P =* χ 0.002), though not in July (X^2^ = 0.11, *P =* 0.74).

**Fig. 3:**
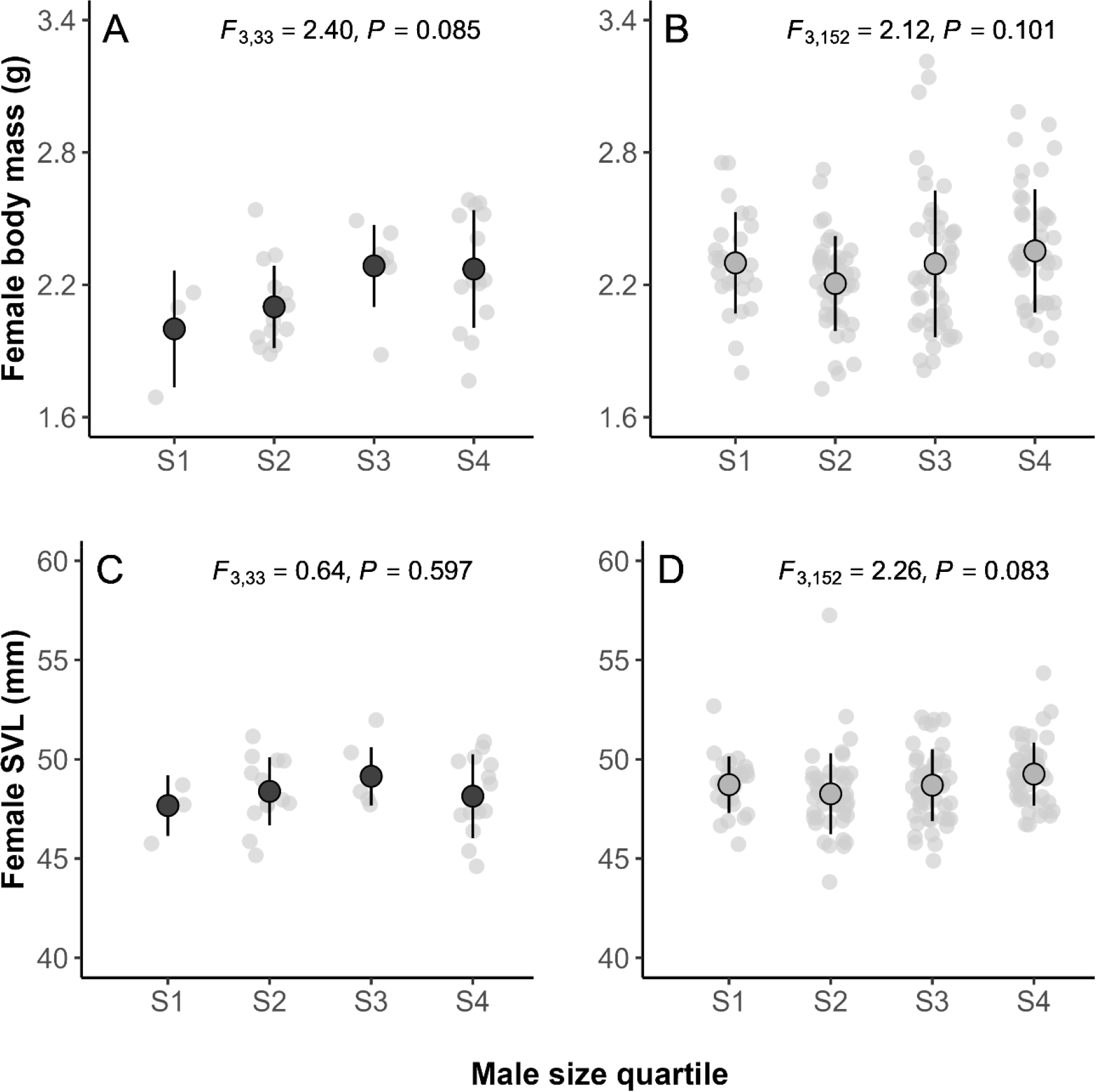
Tests for size-assortative mating with respect to (A-B) body mass or (C-D) snout-vent length (SVL)) of females that mated with males from each size quartile in May (left panels) and in July (right panels), based on the color of fluorescent powder detected on the female. Small filled circles (light grey) are individual values and larger overlaid symbols are mean ± SD values for each quartile. Mating was not strongly size assortative in either May (left) or in July (right), as shown by *F* statistics from a type II ANOVA.

### Comparing behavioral and genetic approaches

Of the 365 males that we successfully genotyped, measured, and included in genetic parentage analysis, 225 were also powdered in either May or July. Total reproductive success estimated from genetic data increased with male size quartile in May (χ^2^ = 39.83, *P* < 0.001, Fig. 4A), although this positive relationship was weaker and not significant in July (χ^2^ = 7.21, *P =* 0.065, Fig. 4B). Combining data across both months confirmed a weak overall relationship between size and reproductive success (Size Quartile: χ^2^ = 6.93, *P* = 0.074), a large effect of month on reproductive success (Month: χ^2^ = 19.30, *P* < 0.001), and a significant difference between months in the relationship between size and reproductive success (Size Quartile x Month: χ^2^ = 14.16, *P* = 0.002, Table 1). Likewise, we found that male mating success increased with size quartile in May (χ^2^ = 32.05, *P* <0.001, Fig. 4C), but this relationship was weaker and not significant in July (χ^2^ = 5.43, *P =* 0.14, Fig. 4D). Pooling data confirmed a significant difference between months in the relationship between size and mating success (Size Quartile: χ^2^ = 5.31, *P* = 0.15, Month: χ^2^ = 14.12, *P* = 0.002, Size Quartile x Month: χ^2^ = 10.52, *P* = 0.014, Table 1). Average mate fecundity was unrelated to male size quartile in May (χ^2^ = 4.01, *P =* 0.26, Fig. 4E) or July (X^2^ = 6.92, *P =* 0.075, Fig. 4F), and pooling data across months revealed a weak but significant tendency for average mate fecundity to decrease with male size (Size Quartile: χ = 9.39, *P* = 0.025, Month: χ = 0.19, *P* = 0.66, Size Quartile x Month: χ = 1.10, *P* = 0.78, Table 1). Competitive fertilization success was unrelated to male size in May (*F_3,62_ =* 1.34, *P =* 0.27, Fig. 4G), in July (*F_3,65_ =* 2.17, *P =* 0.099, Fig. 4H), and when pooling data across months (Size Quartile: *F_3,127_ =* 1.73, *P =* 0.17, Month: *F_1,127_* = 1.87, *P* = 0.35, Size Quartile x Month: *F_3,127_* = 1.82, *P* = 0.15, Table 1).

**Fig. 4:**
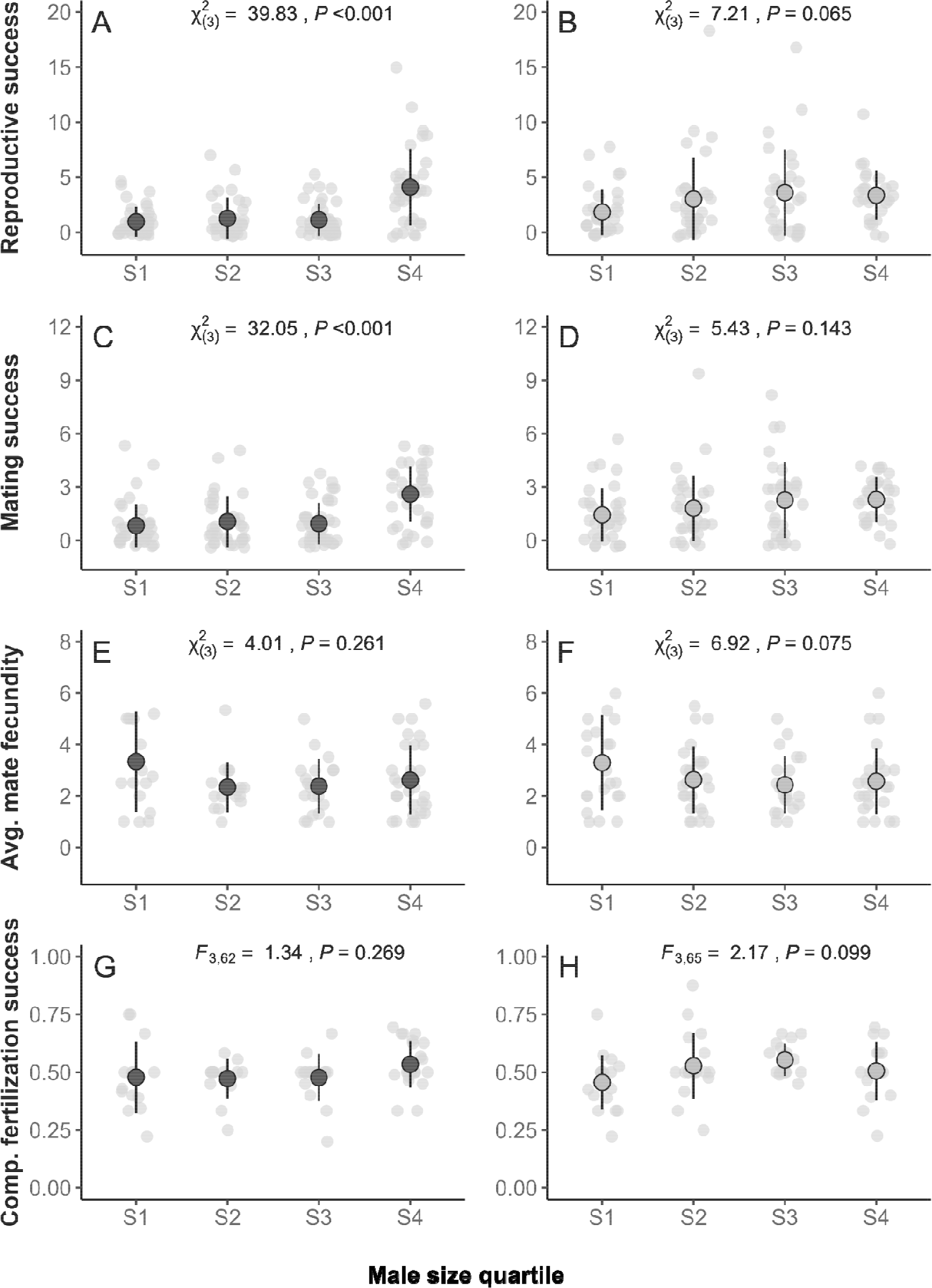
Distribution of (A-B) reproductive success (total number of offspring), (C-D) mating success (total number of mates), (E-F) average mate fecundity (mean fecundity across all mates), and (G-H) competitive fertilization success (mean proportion of offspring sired across all mates, adjusted for the number of competing males), for males powdered in May (left panels) and July (right panels) as a function of their corresponding size quartile. Fitness components were determined using genetic parentage analysis. Small symbols are individual values and larger overlaid symbols are mean ± SD values for each quartile. Large males had significantly higher reproductive success and mating success than small males in May, but not in July. Average mate fecundity and competitive fertilization success did not differ as a function of size quartile in May or in July.

**Table 1:** Summary of coefficient estimates from generalized linear regressions, carried out separately for each measure of fitness as a function of size quartile (ordinal), month (categorical), and their interaction as predictors, when pooling data from May and July: reproductive success (total number of offspring), mating success (total number of mates), average mate fecundity (mean fecundity across all mates), and competitive fertilization success (mean proportion of offspring sired across all mates, adjusted for the number of competing males).

|  | Reproductive<br>success | Mating<br>success | Average mate<br>fecundity | Competitive<br>fertilization<br>success |
| --- | --- | --- | --- | --- |
| *Coefficient* | *Incidence Rate*<br>*Ratios (95% CI)* | *Incidence Rate*<br>*Ratios (95% CI)* | *Incidence Rate*<br>*Ratios (95% CI)* | *$\beta$*<br>*(95% CI)* |
| Intercept | 2.87 ***<br>(2.38 – 3.47) | 1.92 ***<br>(1.63 – 2.25) | 2.76<br>(2.51 – 3.03) | 0.51 ***<br>(0.48 – 0.54) |
| Size Quartile<br>(Linear - L) | 1.56 *<br>(1.06 – 2.30) | 1.44*<br>(1.04 – 2.00) | 0.81*<br>(0.67 – 0.98) | 0.04<br>(-0.02 – 0.09) |
| Size Quartile<br>(Quadratic - Q) | 0.75<br>(0.52 – 1.10) | 0.90<br>(0.65 – 1.23) | 1.12<br>(0.93 – 1.35) | -0.06 *<br>(-0.11 – -0.005) |
| Size Quartile<br>(Cubic - C) | 1.02<br>(0.71 – 1.47) | 0.96<br>(0.70 – 1.30) | 1.00<br>(0.83 – 1.21) | -0.01<br>(-0.06 – 0.05) |
| Month [May] | 0.54 ***<br>(0.41 – 0.71) | 0.63 ***<br>(0.50 – 0.80) | 0.95<br>(0.82 – 1.10) | -0.02<br>(-0.06 – 0.02) |
| Size (L) x Month | 1.64<br>(0.95 – 2.85) | 1.45<br>(0.91 – 2.34) | 1.09<br>(0.83 – 1.44) | -0.00<br>(-0.08 – 0.08) |
| Size (Q) x Month | 2.20 ***<br>(1.27 – 3.82) | 1.64*<br>(1.01 – 2.65) | 1.12<br>(0.83 – 1.50) | 0.09 *<br>(0.01 – 0.17) |
| Size (C) x Month | 1.47<br>(0.85 – 2.56) | 1.46<br>(0.90 – 2.37) | 0.98<br>(0.72 – 1.34) | 0.02<br>(-0.06 – 0.10) |
| n | 262 | 262 | 176 | 135 |
| R<sup>2</sup> | 0.279 | 0.225 | 0.090 | 0.081 / 0.030 |

Estimates indicate the relative increase (Incident Rate Ratio > 1 or $\beta > 0$ ) or decrease (Incident Rate Ratio < 1 or $\beta < 0$ ) with an increase in size quartile for each predictor variable.
Terms are significant if the 95% confidence intervals indicated in brackets do not overlap Incident Rate Ratios at 1 or $\beta$ at 0 (\*\*\* $P < 0.001$ ; \* $P < 0.05$ )

The size distribution of copulation rates inferred from the transfer of fluorescent powder was significantly different from the size distribution of copulation rates estimated from genetic parentage in May (χ = 8.35, *P* = 0.039, Fig. 5A). In particular, males in the second size quartile had more observed copulations than expected from genetic parentage, whereas males in the smallest and largest size quartiles had fewer copulations than expected (Fig. 5A). However, our analyses for May are based on substantially fewer observed copulations (*n* = 37) than our analyses for July (*n* = 156), in which size-specific mating rates observed in the wild did not significantly differ from rates estimated from parentage (χ = 1.41, df = 3, *P =* 0.70, Fig. 5B).

**Fig. 5:**
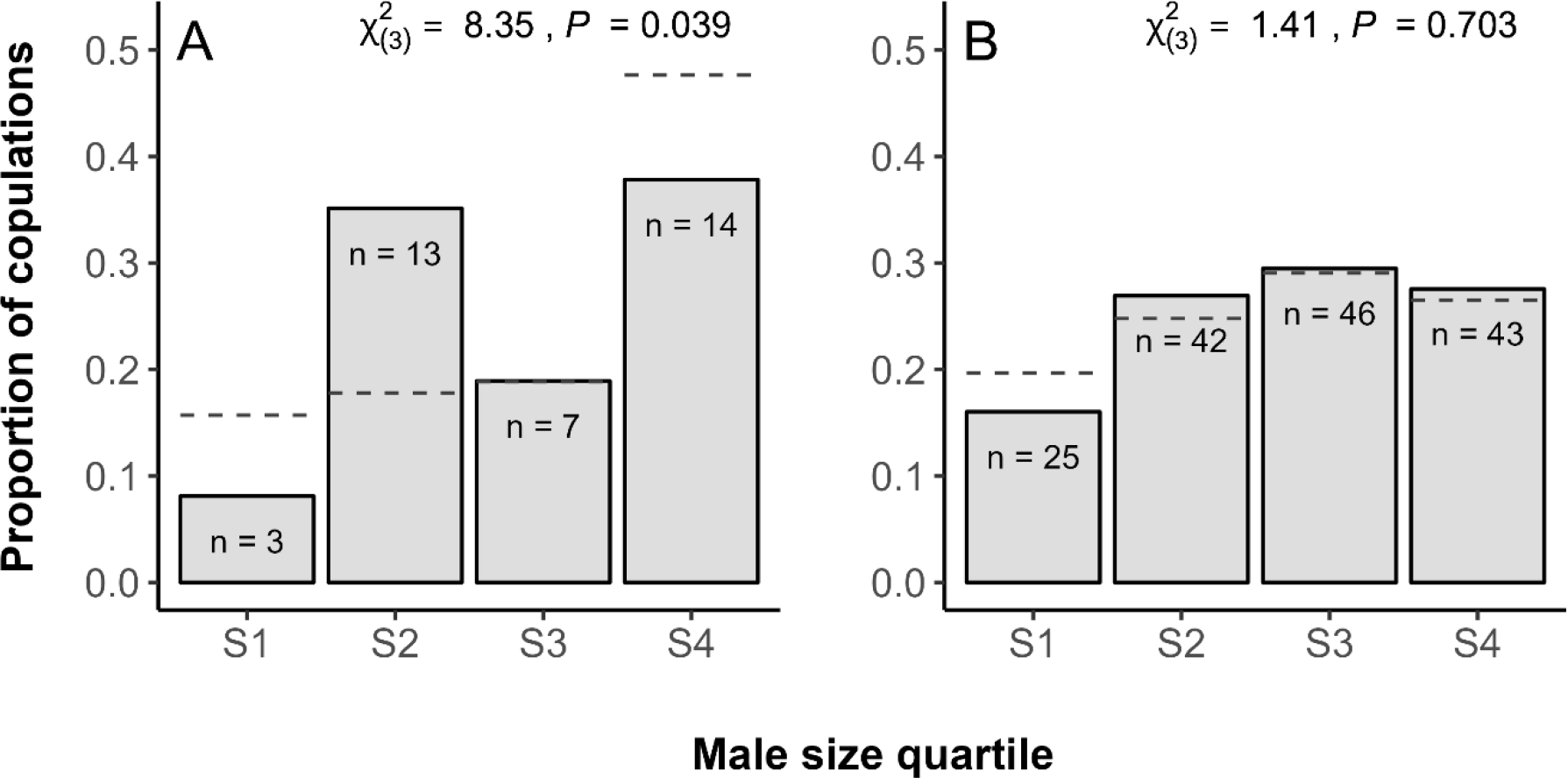
Comparison between behavioral and genetic estimates of male mating success in (A) May and (B) July. Bars represent the proportion of copulations observed for males in each size quartile based on transfer of powder, and dotted black lines indicate the expected proportion of copulations in each size quartile based on the number of mates inferred from genetic parentage data for the same males. Numbers (n) indicate the total number of copulations observed in each size quartile. The observed number of copulations based on powder transfer differed significantly from the expected number based on genetic parentage in May, but not in July.

## Discussion

Pre-copulatory and post-copulatory components of sexual selection can be difficult to disentangle in wild populations, especially for promiscuous species that lack parental care or stable mating pairs. In brown anoles, which lack both, genetic parentage data revealed that 65% of females that produced 2 or more offspring (i.e., females for which multiple paternity could be detected) did so with more than one mate (mean = 1.92, range = 1-4 mates), suggesting the potential for post-copulatory processes to modulate pre-copulatory sexual selection. We detected strong positive directional selection on male body size using estimates of total reproductive success from genetic parentage. Partitioning male reproductive success into its components revealed that the higher reproductive success of larger males was primarily mediated by an increase in their mating success. This result was corroborated by our behavioral assay involving the transfer of fluorescent powder from males to females during copulation, which allowed us to track copulations in the wild and revealed that larger males indeed mated more frequently. By contrast, neither average mate fecundity nor male competitive fertilization success covaried positively with male body size, suggesting that pre-copulatory sexual selection is largely responsible for the strong association between reproductive success and body size in male brown anoles. This was further confirmed by our finding that both behavioral and genetic parentage estimates of mating success were similarly distributed across different male size quartiles. Thus, despite multiple mating by females, post-copulatory processes did not significantly modify pre-copulatory sexual selection for large male body size.

### Body size and mating success

We found that larger body mass is directly associated with greater mating success in the wild (Figs. 1B, 2E-F). This pattern is corroborated by both behavioral and genetic estimates of mating success (Figs. 2E-F, 4C-D). Consequently, larger males sired a greater number of offspring than average throughout the breeding season (Figs. 1A, 4A-B). Our findings are in line with the general consensus that there is strong pre-copulatory sexual selection on male body size in species with extreme male-biased size dimorphism (Stamps et al. 1997; Kingsolver and Pfennig 2004; Fairbairn et al. 2007; Kingsolver and Diamond 2011).

The observed pattern of pre-copulatory sexual selection for large body size is likely due to success in male-male competition (Andersson and Iwasa 1996; Eberhard 1996; Cox et al. 2003; Roff and Fairbairn 2007; Janicke and Fromonteil 2021). Previous studies have shown that larger male anoles are more active (Jenssen et al., 2005; Tokarz, 1985), move across larger areas (Stamps et al. 1997; Kamath and Losos 2018), and are more likely to win in aggressive interactions with other males, resulting in more frequent encounters with females (Steffen and Guyer 2014). This is the case in many other species with male-biased size dimorphism or contest competition (Cox et al. 2003; Fairbairn et al. 2007; Emlen 2008; Janicke et al. 2016; Horne et al. 2020). Although examples of sexual selection via female choice are relatively rare in reptiles (Olsson and Madsen 1995; Tokarz 1995; Cox and Kahrl 2014; Ord et al. 2015; Rosenthal 2017), our study cannot eliminate the role of female choice for large males (Wong and Candolin 2005; Fitze et al. 2008; Karsten et al. 2009; Debelle et al. 2016). Selection due to female choice may occur directly for body size or indirectly through correlated traits such as territory quality, display behaviors, activity levels, and ornaments which signal male aggression and quality (Cooper and Vitt 1993; Censky 1997; Hamilton and Sullivan 2005; Swierk and Langkilde 2013; Flanagan and Bevier 2014; Ord et al. 2015).

Although genetic estimates of reproductive success and mating success were strongly correlated with body size or size quartiles measured early in the breeding season (May, Figs. 1A- B; 4A, 4C), they were not strongly correlated with size quartiles measured later in the breeding season (July, Figs. 4B, 4D; Table 1). Estimates of mating success from both behavioral and genetic measures were similarly high for males beyond the first size quartile in July (Figs. 2F; 4D). This may indicate that, beyond a certain threshold, the advantage of large size in agonistic interactions with other males can saturate (Cox and Calsbeek 2010a; Reedy et al. 2017).

### Body size and average mate fecundity

Male body mass was mostly unrelated to, or sometimes even negatively correlated with, the average fecundity of female partners (Figs. 1C, 4E-F; Table 1). This may reflect the fact that the relationship between female body mass and male size quartile was weak and nonsignificant within each month (Fig. 3A-B), and female mass itself was unrelated to fecundity. Although an alternative measure of female size (SVL) was significantly related to fecundity, consistent with previous work showing that larger female anoles may achieve a higher reproductive output (Warner and Lovern 2014; Duryea et al. 2016) by laying eggs more frequently (Cox and Calsbeek 2011), we did not find any association between male size quartile and female SVL (Fig. 3C-D). Thus, neither body mass nor SVL of females provided a strong intermediate linking male size to female fecundity via size-assortative mating. These findings are consistent with the general observation that size-assortative mating is rare, particularly in species with male-biased sexual size dimorphism, such as anoles (Shine et al. 2001; Hofmann and Henle 2006; Harrison 2013; Rios Moura et al. 2021). When mate choice has been detected in anoles, males appear to prefer novel females rather than larger females (Tokarz 1992; Orrell and Jenssen 2002). This would be expected if males are primarily under selection to mate with a greater number of females, rather than more fecund females. In contrast, larger males often mate with larger and/or more fecund females in species with female-biased sexual size dimorphism (Verrell 1989; Olsson 1993; Whiting and Bateman 1999; Cox et al. 2005; John-Alder et al. 2009; Jiang et al. 2013).

### Body size and competitive fertilization success

Consistent with previous findings in brown anoles, we found that over 50% of the females having at least two genotyped offspring produced these offspring with more than one mate (Calsbeek et al. 2007; Duryea et al. 2016; Kahrl et al. 2021). Moreover, at least 6% of the females in our powdering studies mated with multiple partners within a short 2-5 day span. The actual frequency of multiple mating is likely to be much higher because our powdering method cannot detect instances of multiple mating within size quartiles, and because our ability to detect multiple paternity is limited by the relatively low number of offspring produced by females (mean = 2.06, range = 1-9 offspring). Although multiple mating by females was common, in situations where females produced offspring with multiple males, male size was unrelated to fertilization success (Figs. 1D, 4G-H, Table 1).

Post-copulatory processes can oppose pre-copulatory selection on a given trait if investment in corresponding fitness components is drawn from the same limited resource, or if the genetic covariance among fitness components is negative (Roff and Fairbairn 2007; Parker et al. 2013). Accordingly, inter- and intraspecific comparisons across several lineages, including reptiles, have shown that traits typically subjected to pre-copulatory selection trade-off with those under post-copulatory selection (Moczek and Nijhout 2004; Fitzpatrick et al. 2012; Dines et al. 2015; Kahrl et al. 2016; Somjee et al. 2018). On the other hand, when there is high mean and variance in resource acquisition, this association is likely to be positive since any increase in resource availability allows for more investment in both pre- and post-copulatory competition, (Saeki et al. 2014; Simmons et al. 2017). Consistent with this idea, several intra-specific studies have reported a positive correlation between targets of pre-copulatory sexual selection and ejaculate traits (*reviewed in* Mautz et al. 2013; Supriya et al. 2019). Although some studies report positive associations between standardized fertilization success and traits such as body size, singing effort and/or weapon size (Preston et al. 2001; Hosken et al. 2008; Turnell and Shaw 2015; House et al. 2016), others report negative associations (Danielsson 2001; Evans et al. 2003; Kelly and Jennions 2011). However, our findings are consistent with those studies in which male fertilization success is unrelated to body size or ornament size (Keogh et al. 2013; Rose et al. 2013; Flanagan et al. 2014; McDonald et al. 2017). This may indicate that investment in mate acquisition does not trade off with investment in fertilization success, possibly due to the predicted low cost of producing ejaculates when these are distributed across several matings (Hayward and Gillooly 2011; Parker 2016; Kahrl et al. 2021; *but see* Kahrl and Cox 2015).

One caveat is that our measure of competitive fertilization success required us to exclude all instances in which a single male sired all of the offspring produced by a female, potentially excluding extremely strong or weak sperm competitors from our analysis (Fig. S2D, S3B).

However, failure to account for the number of competing males in this way may result in spurious correlations. This is because the estimated proportion of offspring sired by a male will increase, regardless of the focal male’s competitive ability, if females produce offspring with fewer mates (Rose et al. 2013; Devigili et al. 2015; McCullough et al. 2018). Indeed, when we used unadjusted fertilization success in brown anoles, we found significant, albeit very weak, positive selection on male body size (Fig. S3). Thus, post-copulatory selection on body size may be weaker in natural populations than previously reported by studies using unadjusted measures of male fertilization success (Preston et al. 2001; Hosken et al. 2008; Turnell and Shaw 2015; House et al. 2016). Our study suggests that, at least for body size, post-copulatory selection is negligible compared to pre-copulatory selection. It is more likely that post-copulatory selection acts primarily on male ejaculate traits, as has been demonstrated in brown anoles (Kahrl and Cox 2015), and that it may operate independent of male body size (Kahrl et al. 2021).

### Comparing behavioral and genetic measures of mating success

We found a close association between measures of size-specific mating success derived from genetic parentage and those inferred from copulations in the field, particularly in July (Fig. 5B). This highlights the utility of fluorescent powder transfer as a relatively inexpensive and effective method for detecting copulations, particularly in natural populations, and for linking mating success to broad categories of phenotypic variance. Our findings are in line with other studies that have found behavioral proxies, such as the frequency of male-female associations in space and time, to be closely predictive of the realized mating and reproductive success of males (Kamath and Losos 2018; Olsson et al. 2019; Baird and York 2021). However, our technique is much easier to execute compared to detailed observations of individual copulations or movements, at least in our focal species. Thus, it can be used to uncover associations between mating success and categorical simplifications of continuous traits (as in this study), naturally categorical traits or groups (e.g., morphs), or experimental treatments (e.g., Wittman et al. 2022). It can also be used to uncover mating patterns of secretive or spatially dispersed species that can be difficult to observe in the wild for long hours (Gosden and Svensson 2007; Johnson et al. 2014). Nonetheless, behavioral estimates of size-specific mating success based on powder transfer only corresponded closely with genetic mating success when extensive sampling of the female population was possible and when mating rate was high (Fig. 5A-B). For example, in May, we only sampled females for 1 day and the inferred mating rate was half of that seen in July, when we sampled for 5 days (Fig. 5A-B). Perhaps as a result, the relatively low number of observed copulations in May differed significantly from our expected distribution of size-specific mating success, which was likely more accurate because it was based on a much larger number of inferred copulations from genetic parentage (Fig. 5A). Thus, behavioral observations or genetic parentage alone may not adequately capture fitness when populations are partially sampled or if mating is infrequent within a short sampling period.

## Conclusions

Overall, our study confirms that large body size is associated with higher reproductive success in brown anoles, and that this is primarily due to the increased mating success of large males. Although previous work has suggested that sexually antagonistic viability selection may favor large male size and promote male-biased sexual size dimorphism in this species (Cox and Calsbeek 2010a; *but see* Cox and Calsbeek 2015), our results support a parallel body of recent work suggesting that sexual selection also strongly favors large male size (Tokarz 1985; Jenssen et al. 2005; Duryea et al. 2016; Kamath and Losos 2018). Importantly, we extend this work by specifically resolving the importance of pre-copulatory sexual selection and linking large male size to both behavioral and genetic measures of mating success. Our results further illustrate that strong pre-copulatory sexual selection and extremely male-biased sexual size dimorphism can occur even in promiscuous mating systems in which access to females cannot be monopolized and multiple paternity is common. Finally, our findings emphasize the importance of incorporating both behavioral and genetic methods in the same study to achieve a more robust understanding of the roles of pre- and post-copulatory processes in sexual selection.

## Data Availability

All code and data needed to reproduce the results presented in this paper will be available on Dryad Digital Repository at https://doi.org/10.5061/dryad.c866t1gbb

## Acknowledgements

We thank D. Warner for logistical support during field collections and genotyping, T. Carlson, C. Giordano, B. Ferrer and B. Beck for assistance during field sampling and pilot studies, E. Bruns, L. Galloway and T. J. Chechi for lending DayGlo fluorescent powders, N. Campbell and GTSeek LLC for advice on genotyping and library preparation of samples, and M. Hale, P.L.H. de Mello, and D. Nondorf for their input on earlier drafts of the manuscript. This study was conducted under permits from the Guana Tolomato Matanzas National Estuarine Research Reserve and approval of the University of Virginia Animal Care and Use Committee (protocol 3896).

## Supplementary Figures

**Fig. S1.**
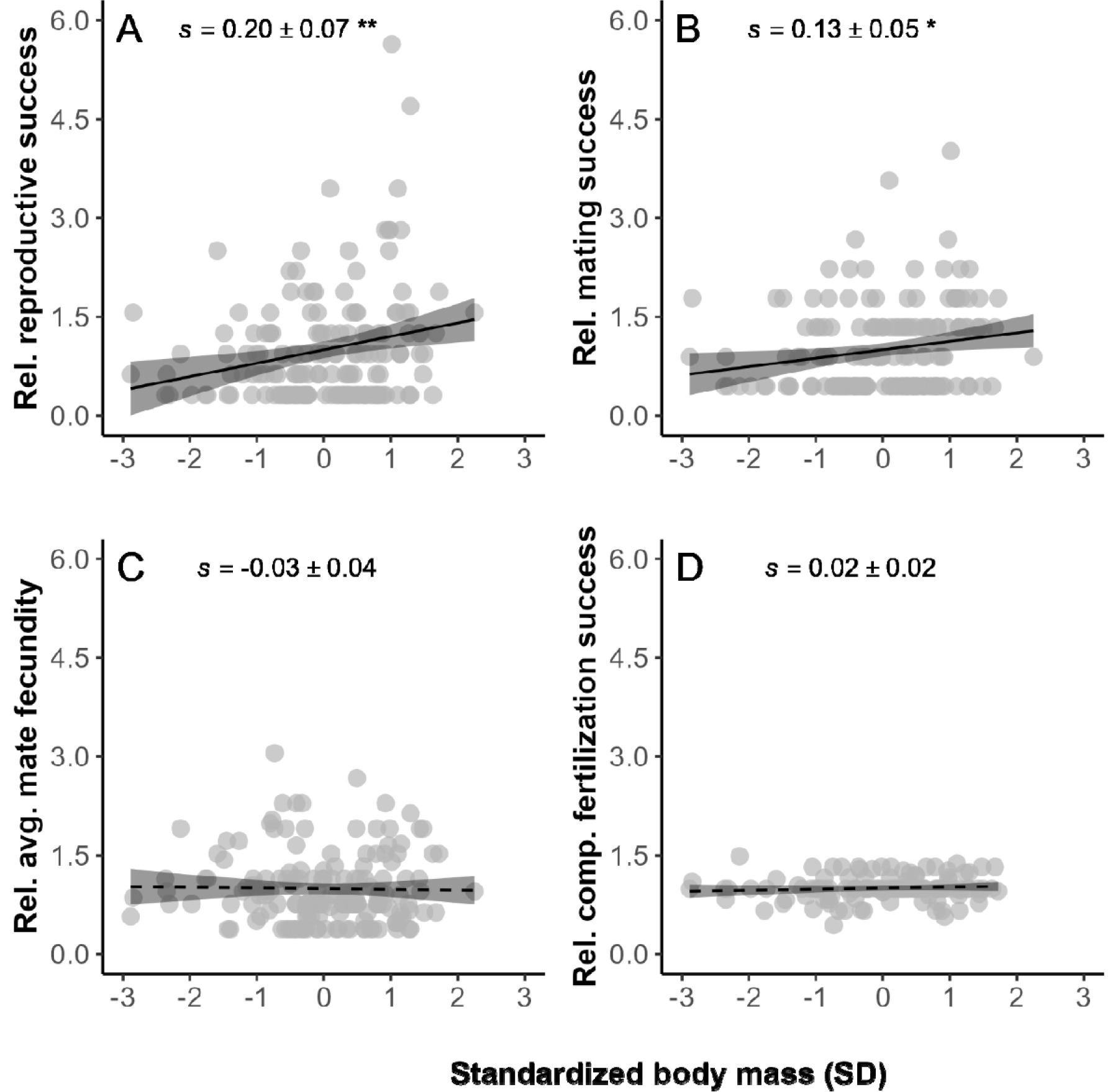
Linear selection on standardized adult body mass as a function of different components of male fitness, including relative measures of (A) reproductive success (total number of offspring), (B) mating success (total number of mates), (C) average mate fecundity (mean fecundity across all mates), and (D) competitive fertilization success (mean proportion of offspring sired across all mates, adjusted for the number of competing males). Here, individuals that were not assigned offspring in parentage analysis have been excluded from all regressions to avoid inflating the relative contribution of mating success to total reproductive success by assuming that these males failed to mate. Trendlines (with 95% CI) from linear regressions are used to visualize selection. Solid lines and asterisks in panels A-B indicate significant selection differentials (*s*) while the dotted lines indicate non-significant selection differentials (** *P* < 0.01; * *P* < 0.05). Excluding males without any assigned progeny reduces the overall strength of selection on size by 50% (from *s* = 0.40 to 0.20, compare with Figs. 1 or S3), but only slightly reduces the proportion of that selection attributable to variance in mating success (from *s* = 0.33 or 82.5% to *s* = 0.13 or 65%, compare with Figs. 1 or S2).

**Fig. S2.**
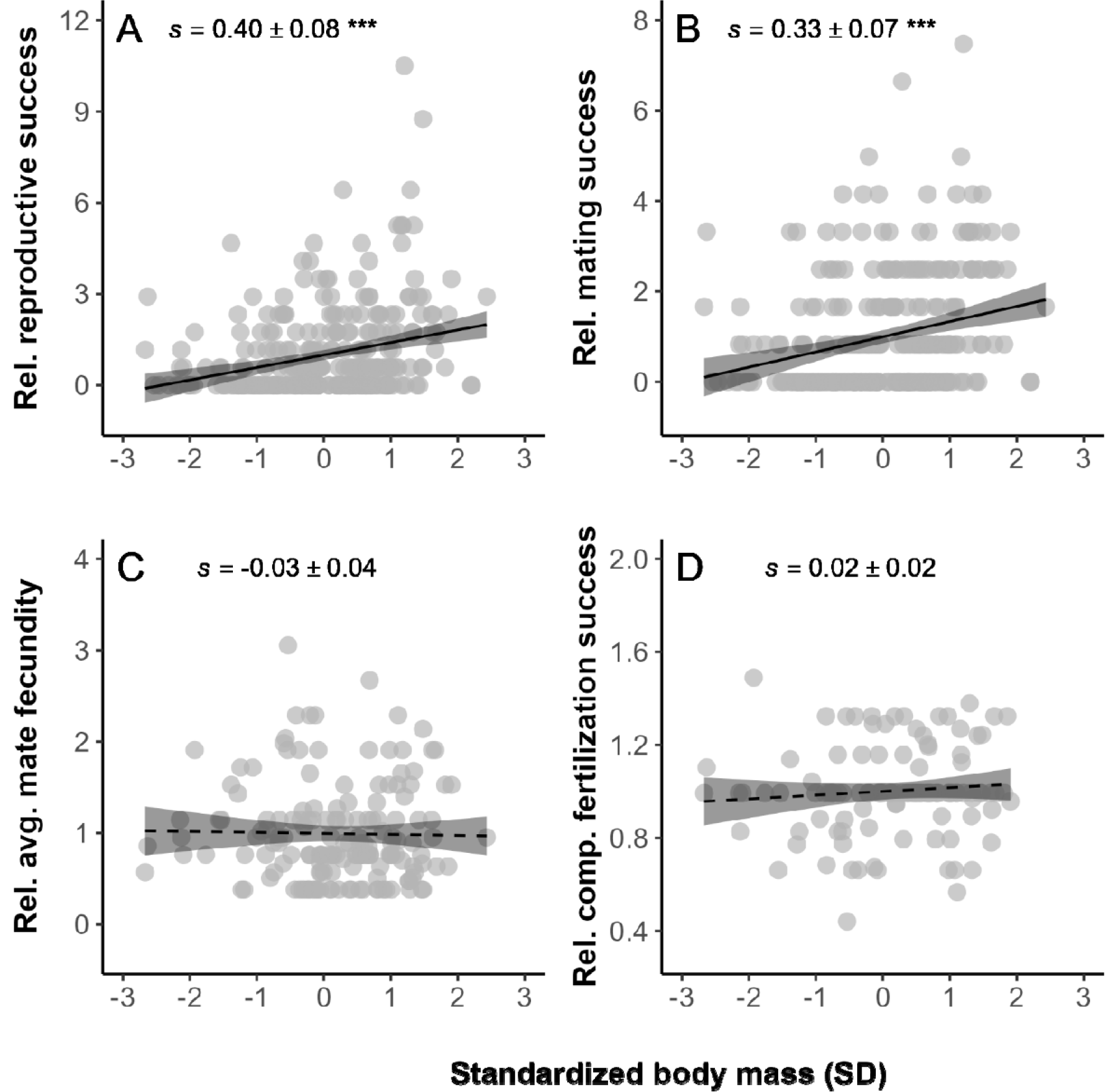
Linear selection on standardized adult body mass as a function of different components of male fitness, including relative measures of (A) reproductive success (total number of offspring), (B) mating success (total number of mates), (C) average mate fecundity (mean fecundity across all mates), and (D) competitive fertilization success (mean proportion of offspring sired across all mates, adjusted for the number of competing males). Here, the y-axis has been scaled to include only the range of data points for each fitness component to clearly visualize the selection differentials estimated in Fig 1. Trendlines (with 95% CI) from linear regressions are used to visualize selection. Solid lines and asterisks in plots A-B indicate significant selection differentials (*** *P* < 0.001) while the dotted lines indicate non-significant selection differentials (*P* > 0.05).

**Fig. S3.**
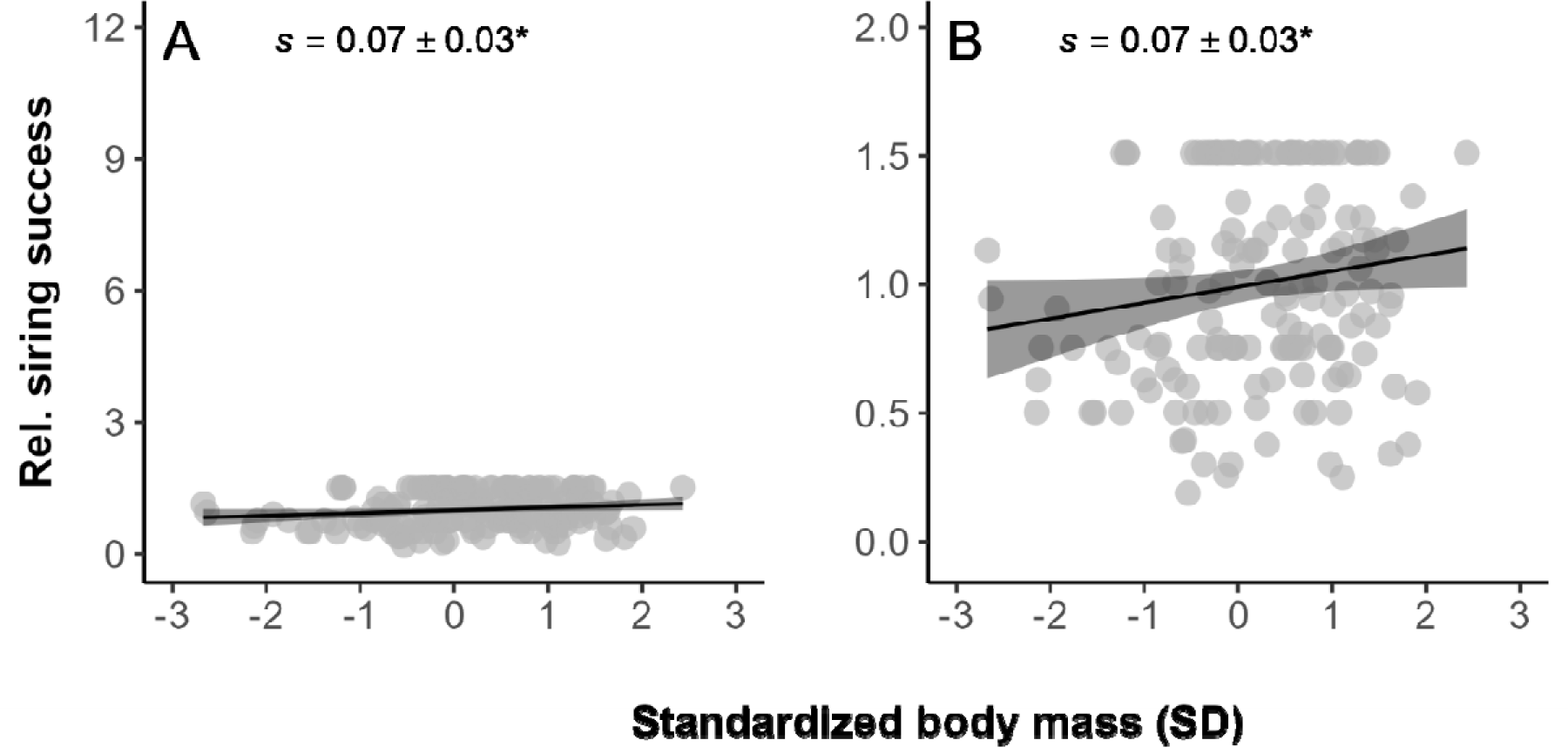
Linear selection on standardized adult body mass as a function of relative siring success of males (not adjusted for competitor males) (A) along the same scale as the y-axis in Fig 1 or (B) with y-axis scaled to include only the range of data points for the fitness component to clearly visualize the same selection differential. Trendlines (with 95% CI) from linear regressions are used to visualize selection. Solid lines and asterisks indicate significant selection differentials (* *P* < 0.05). Selection due to unadjusted measures of fertilization success on male body mass was significant and three times stronger than that due to competitive fertilization success (from s = 0.02 to 0.07, compare with Figs. 1D or S2D)

